# Tracing LYVE1^+^ peritoneal fluid macrophages unveils two paths to resident macrophage repopulation with differing reliance on monocytes

**DOI:** 10.1101/2025.03.19.644175

**Authors:** Alexandre Gallerand, Jichang Han, Rachel L. Mintz, Jing Chen, Daniel D. Lee, Mandy M. Chan, Tyler T. Harmon, Xue Lin, Christopher G. Huckstep, Siling Du, Tiantian Liu, Jonathan Kipnis, Kory J. Lavine, Joel D. Schilling, S. Celeste Morley, Bernd H. Zinselmeyer, Kenneth M. Murphy, Gwendalyn J. Randolph

## Abstract

Mouse resident peritoneal macrophages, called large cavity macrophages (LCM), arise from embryonic progenitors that proliferate as mature, CD73^+^Gata6^+^ tissue-specialized macrophages. After injury from irradiation or inflammation, monocytes are thought to replenish CD73^+^Gata6^+^ LCMs through a CD73^-^LYVE1^+^ LCM intermediate. Here, we show that CD73^-^LYVE1^+^ LCMs indeed yield Gata6^+^CD73^+^ LCMs through integrin-mediated interactions with mesothelial surfaces. CD73^-^LYVE1^+^ LCM repopulation of the peritoneum was reliant upon and quantitatively proportional to recruited monocytes. Unexpectedly, fate mapping indicated that only ∼10% of Gata6-dependent LCMs that repopulated the peritoneum after injury depended on the LYVE1^+^ LCM stage. Further supporting nonoverlapping lifecycles of CD73^-^LYVE1^+^ and CD73^+^Gata6^+^ LCMs, in mice bearing a paucity of monocytes, Gata6^+^CD73^+^ LCMs rebounded after ablative irradiation substantially more efficiently than their presumed LYVE1^+^ or CD73^-^ LCM upstream precursors. Thus, after inflammatory insult, two temporally parallel pathways, each generating distinct differentiation intermediates with varying dependencies on monocytes, contribute to the replenish hment of Gata6^+^ resident peritoneal macrophages.

## INTRODUCTION

Resident macrophages are found at high concentrations in the peritoneal fluid to promote host defense and tissue homeostasis in the visceral body cavity (*1–3*). The major resident peritoneal macrophages arise from embryonic progenitors capable of sustained self-renewal (*4–7*). These resident macrophages are called large peritoneal macrophages (*8*), and they coexist with small peritoneal macrophages (*9*) that arise from monocytes (*10*). Because the pleural and pericardial serosal cavities have similarly distinct tissue resident macrophages as the peritoneal cavity (*11–13*), more universal terms for these macrophages in the three serosal tissue compartments are ‘large cavity macrophages’ (LCMs) and ‘small cavity macrophages’ (SCMs). LCMs have a unique identity in the mouse, readily identified by their surface expression of ICAM2 (CD102), CD62P (P-selectin), and CD73 that other tissue resident macrophages do not express (*14–17*).

Among immune cells, the transcription factor Gata6 is uniquely expressed in LCMs, and this transcription factor regulates LCM abundance (*15–17*) and expression of enriched LCM genes like CD73 and CD62P (*15, 18*). However, a few LCMs persist in mice whose macrophages have deleted *Gata6*. These Gata6-deficient macrophages retain certain unique features of LCMs, including surface expression of ICAM2 (*15, 16*), and while they no longer express CD73 and CD62P, they co-express LYVE1 and CD206 (*16, 18*). Explanations for the reduced numbers of LCMs in the absence of Gata6 include a higher propensity for cell death (*16*), impaired proliferation (*17*), and entrapment of differentiation intermediates in the nearby omentum (*15*), a mesothelial cell-lined specialized peritoneal fold containing adipose tissue rich in lymphoid follicles that is attached to the greater curvature of the stomach. Indeed, the identification of large numbers of apparent ICAM2^+^ LCM differentiation intermediates that were not yet Gata6^+^ in the omentum prompted Okabe and Medzhitov to suggest that the omentum provides essential signals, including the production of retinoic acid, that support the induction of Gata6 and final maturation of LCMs (*15*). This interesting concept positioned the omentum as a site not just for macrophage clearance during the resolution of inflammation (*1, 19*) but also for their developmental origin. Undefined omental factor(s) and retinoic acid were proposed as niche factors that supported LCM development and maintenance (*15*). LCMs indeed rely on receipt of retinoic acid signals for maintenance (*12, 15, 20, 21*), in particular upon expression of RXRα and RARγ (*20, 21*). Retinoic acid is generated in the omentum by specialized fibroblasts, but deleting the capacity of these fibroblasts to produce retinoic acid did not impact the accumulation of resident peritoneal macrophages in peritoneal fluid (*22*), underscoring the expectation that retinoic acid might arise from sources beyond the omentum. One alternative source may be Wt1^+^ mesothelial cells that are more broadly distributed along the surface lining of the peritoneal cavity (*12*). Mesothelial cell-produced mesothelin and Muc16 may influence Gata6 expression in LCMs (*23*). Additionally, the mesothelium is an important source of macrophage colony-stimulating factor 1 (CSF1) that drives macrophage proliferation and survival (*24*).

In homeostasis, most LCMs remain of embryonic origin, although monocytes slowly but steadily contribute to tissue resident macrophages over time (*25–28*). Monocyte contribution to the Gata6^+^ resident peritoneal macrophage pool accelerates during sterile inflammation or parasitic infection (*11, 16, 18, 29*). Repopulation of LCMs after irradiation-induced genotoxic stress arising in the context of bone marrow transplant (BMT) is entirely dependent upon bone marrow-derived cells (*7*) that are thought to arise from CCR2^+^ monocytes (*25, 26*). Indeed, the extent to which recruited monocytes repopulate CD73⁺CD62P⁺ LCMs following helminth infection varies across inbred mouse strains, affecting their immune response and physiological recovery (*11, 16, 18, 29, 30*).

Monocytes that enter the peritoneal cavity transition first to ICAM2^+^ LCMs that resemble the LYVE1^+^ macrophages earlier described in Gata6^ΔMac^ mice (*16, 31*), before becoming Gata6^+^ LCMs (*11, 16, 18, 25, 29, 30*). These so-called “converting macrophage” differentiation intermediates are characterized not only by the induction of ICAM2 but also by the expression of CD206, LYVE1, and folate receptor-β (*11, 16, 18*). Passage through this intermediate stage of differentiation appears to give way to the expression of Gata6, Tim4, and Gata6-regulated CD73 and CD62P (*15, 18*). This maturation is accompanied by augmented F4/80, retention of ICAM2, and loss of CD206 and LYVE1 expression (*11, 18, 25, 31*).

In this study, we sought to elucidate the details defining the stages of LCM differentiation and the involvement of peritoneal lining tissues like the omentum and mesentery in this differentiation. We show that the adhesion of LCMs to peritoneal linings influences the differentiation of the LYVE1^+^ LCM ‘converting macrophages’ and that LYVE1^+^ LCMs could differentiate to CD73^+^CD62P^+^ LCMs in a lineage tracing model. However, quite unexpectedly, our findings failed to show the anticipated result that Gata6-dependent LCMs obligatorily pass through a LYVE1^+^ ‘converting macrophage’ differentiation intermediate after a BMT that enforces LCM turnover. Instead, the data revealed that differentiation of LCMs bifurcates during LCM replenishment from bone marrow precursors, such that the LYVE1^+^ LCM trajectory existed but was a minor trajectory in the C57Bl/6 mouse. This trajectory was, however, quantitatively dependent upon monocytes in the steady state or after BMT, as was determined in mice wherein Zeb2 expression in monocytes is impaired such that monocyte development is blocked (*32, 33*). Although all Gata6^+^ LCMs relied on CCR2 and arose from Ms4a3^Cre^-labeled precursors (*26*), suggesting a relationship to monocyte-like cells, most did not induce LYVE1 during differentiation. Indeed, CD73^+^Gata6^+^ LCMs showed limited quantitative reliance on monocytes, as the repopulation of most LCMs was scarcely affected in Zeb2 mutant mice lacking monocytes. These data reveal unexpectedly that LCMs in the peritoneal cavity develop along two parallel pathways with different intermediates, dependence on interactions with peritoneal surface tissues, and reliance on monocytes. Our previous studies characterizing human LCMs of the peritoneal cavity (*31*) raise the possibility that only one of these two pathways is recapitulated in humans.

## RESULTS

### Peritoneal surface and fluid distribution of ICAM2^+^ LCMs at different differentiation stages

LYVE1 is strongly expressed by peritoneal converting macrophages (*18*). It is also expressed by mesothelium-associated macrophages (*34*), omentum-associated macrophages (*35*), and Gata6-deficient LCMs (*16*) that have been reported to accumulate in the omentum (*15*). However, only some of these macrophages would be expected to fall within the LCM pool defined by the surface expression of ICAM2. We set out to compare the distribution of ICAM2^+^LYVE1^+^ macrophages, along with other macrophages, in peritoneal fluid compared to the omentum and to further understand their phenotypes. Because isolating single cells from the omentum requires enzymatic digestion, whereas isolating fluid-residing cells does not, we developed a method to account for the effects of enzymatic digestion on surface marker expression (**Fig. 1A**). Omentum samples were isolated from CD45.1 congenic C57BL/6 mice, minced and mixed with peritoneal fluid from either WT or Lyz2cre x *Gata6*^fl/fl^ (Lyz2^ΔGata6^) CD45.2 C57BL/6 mice (**Fig. 1A**). Samples were then digested with collagenase IV, surface stained, and analyzed using spectral flow cytometry. Samples resulting from mixes of individual biological replicates were concatenated prior to unsupervised analysis (**Fig. 1B**). Doublets and dead cells were excluded before gating on F4/80^+^ CD11b^+^ MerTK^+^ macrophages. Then, dimensionality reduction and cell clustering were conducted, allowing the identification of 8 clusters (**Fig. 1B-D**). Cluster 4 was most abundant in the WT peritoneal fluid sample, while it was almost absent in the Lyz2^ΔGata6^ peritoneal fluid (**Fig. 1C**). Conversely, cluster 3 was rare in the WT fluid sample but predominated in the Lyz2^ΔGata6^ fluid (**Fig. 1C-D**). Cluster 7 was only present in peritoneal fluid samples and accumulated in Lyz2^ΔGata6^ mice (**Fig. 1C**). Clusters 1 and 2 defined most omental macrophages (**Fig. 1C-D**). Cluster 0 was exclusively associated with the omentum, where it was defined by expression of CD169, as previously described (*36*) (**Fig. 1D**). Clusters 2, 3, and 5 were present in both omental and peritoneal fluid samples. ICAM2 expression could be detected in clusters 2, 3, and 4 (**Fig. 1C-E**). Cluster 4 also displayed expression of the Gata6-associated genes CD73 and CD62P (*14, 15*) (**Fig. 1E**). After manually gating on CD73^+^CD62P^+^ LCMs (**fig. S1A**), we confirmed that this population highly expressed Gata6, measured by intracellular staining, compared to CD73^-^CD62P^-^ICAM2^+^ LCMs in which Gata6 was absent and resembled that of LCMs engineered to delete *Gata6* (**fig. S1B**). Thus, the surface expression of CD73 and CD62P served as faithful surrogates for Gata6 expression. Macrophages in clusters 2 and 3 were characterized by a lower expression of F4/80 and a higher expression of CD206, folate receptor β (FRβ), and LYVE1 compared to those in cluster 4 (**Fig. 1C, E, G-I**). This expression pattern aligns with the signature of macrophages undergoing conversion into mature peritoneal cavity macrophages, as previously described (*18*). The peritoneal wash from Lyz2^ΔGata6^ mice was enriched in ICAM2^+^LYVE1^+^ macrophages that also expressed CD206 and folate receptor β (*16*) but lacked CD73, defined by clusters 2 and 3 (**Fig. 1E, G-I**).

**Figure 1.**
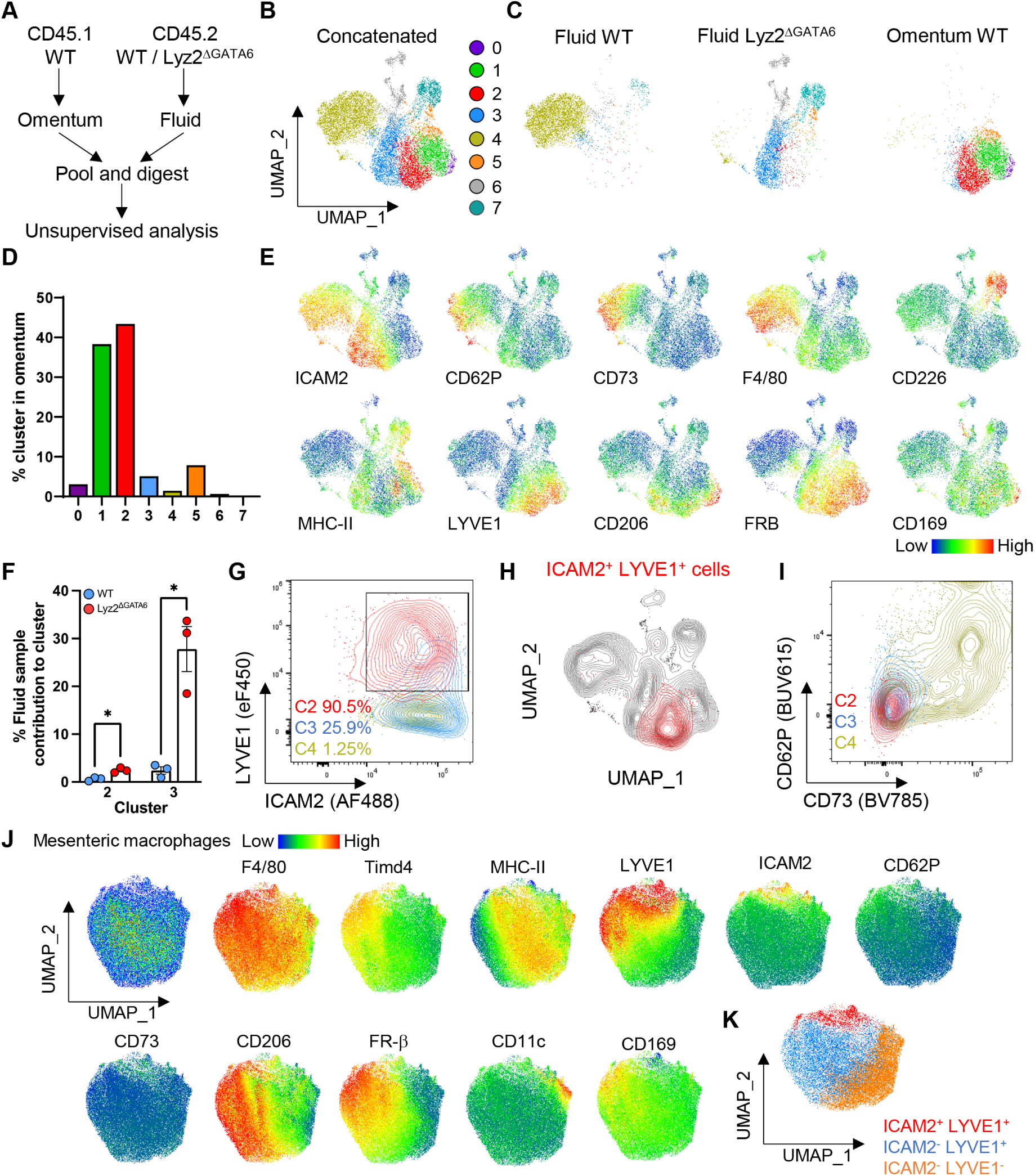
Phenotyping and comparison of ICAM2^+^ macrophages from the omentum and peritoneal fluid. (**A**) Experimental scheme used to acquire data for subsequent unsupervised analysis. CD45.2^+^ cells retrieved from the peritoneal wash of Gata6^fl/fl^ (WT) or Lyz2^ΔGata6^ mice were added to the minced CD45.1 omental tissue, and cells were digested at 37°C for 30 min. (**B, C**) UMAPs showing unsupervised clustering of omental and peritoneal wash macrophages analyzed via flow cytometry. (**D**) Proportions of each cluster’s contribution arising from the omentum. (**E**) UMAP plots representing the expression of macrophage markers used to define cluster identity. (**F**) Proportions of WT and Lyz2^ΔGata6^ cells found among clusters 2 and 3. (**G**) Flow cytometry plot showing expression of LYVE1 and ICAM2 in macrophages from clusters 2, 3, and 4. (**H**) UMAP showing the distribution of ICAM2^+^ LYVE1^+^ macrophages (red) compared with all other macrophages (gray) contained in the dataset. (**I**) Flow cytometry plot showing expression of CD62P and CD73 in macrophages from clusters 2, 3, and 4. (**J**) UMAPs displaying expression of the indicated surface markers by mesenteric macrophages. (**K**) Overlay of manually gated mesenteric macrophage populations. Panels A-I contain concatenated data from CD45.1 mice (n=3), CD45.2 WT mice (n=3), CD45.2 Lyz2^ΔGata6^ mice (n=3) and represent three similar experiments. Panels J-K are representative of two independent experiments. An ANOVA was used for statistical analysis in panel F. See also Extended Data Figure 1. *P<0.05.

Clusters 2 and 3 were distinguished by the site where they predominantly localized: cells from cluster 2 overwhelmingly resided in the omentum, while cells from cluster 3 overwhelmingly resided in peritoneal fluid (**Fig. 1D**). When we probed further to define phenotypic differences between these two ICAM2^+^ clusters, we observed that cluster 2 uniformly expressed LYVE1 while only a fraction of cluster 3 and almost none of cluster 4 did so (**Fig. 1G**). Consistent with this observation, total ICAM2^+^ LYVE1^+^ macrophages from the concatenated dataset localized mainly in cluster 2 (**Fig. 1H**). These data were consistent with the possibility that cluster 2 in the omentum corresponds specifically to a differentiation stage referred to as ‘converting macrophage’ intermediates (*18*), which precedes the development of macrophages in clusters 3 and 4 that appear to be in the process of downregulating LYVE1 while acquiring CD73 and CD62P expression. The data imply that LCM differentiation and maturation involves anatomic repositioning, with cells transitioning from peritoneal surfaces like the omentum into the peritoneal fluid as they progress through differentiation intermediates.

We also analyzed whether these populations were present in mesenteric tissue. The majority of LYVE1^+^ macrophages of the mesentery were ICAM2^-^ macrophages, likely accounted for by those previously described to reside in mesothelial compartments (*34*) and the omentum (*36*), including many expressing CD169 (*36*) (**Fig. 1J**). A distinct subset of LYVE1^+^ICAM2^+^ macrophages was also associated with the mesentery (**Fig. 1J-K**). Collectively, these data suggested that LYVE1^+^ICAM2^+^ macrophages from the omentum or mesentery might mature into LCMs found in peritoneal fluid, fitting with the paradigm that peritoneal surface tissues serve as a seeding ground for the development of mature LCMs (*15*).

### Peritoneal surfaces harbor LYVE1^+^ LCMs that later differentiate into peritoneal fluid CD73^+^CD62P^+^ LCMs

Supporting the concept of peritoneal surfaces as reservoirs for LYVE1^+^ICAM2^+^ LCMs, brief trypsin exposure detached LYVE1^+^ LCMs from the mesothelial surfaces of peritoneal organs after an initial in vivo peritoneal lavage without trypsin removed freely floating cells. Flow cytometry analysis confirmed that CD11b^+^ICAM2^+^CD73^-^LYVE1^+^ macrophages were highly enriched in the brief trypsin digest (surface) compared with the population retrieved in the peritoneal lavage (∼35% versus <1%) (**Fig. 2A**). More than 60% of ICAM2^+^ macrophages from the digested omentum and about 40% of ICAM2^+^ macrophages digested from the mesenteries were LYVE1^+^ and all were CD73^-^ (**Fig. 2B**). In contrast, CD73^+^ cells were only recovered abundantly in the peritoneal fluid (**Fig. 2A**).

**Figure 2.**
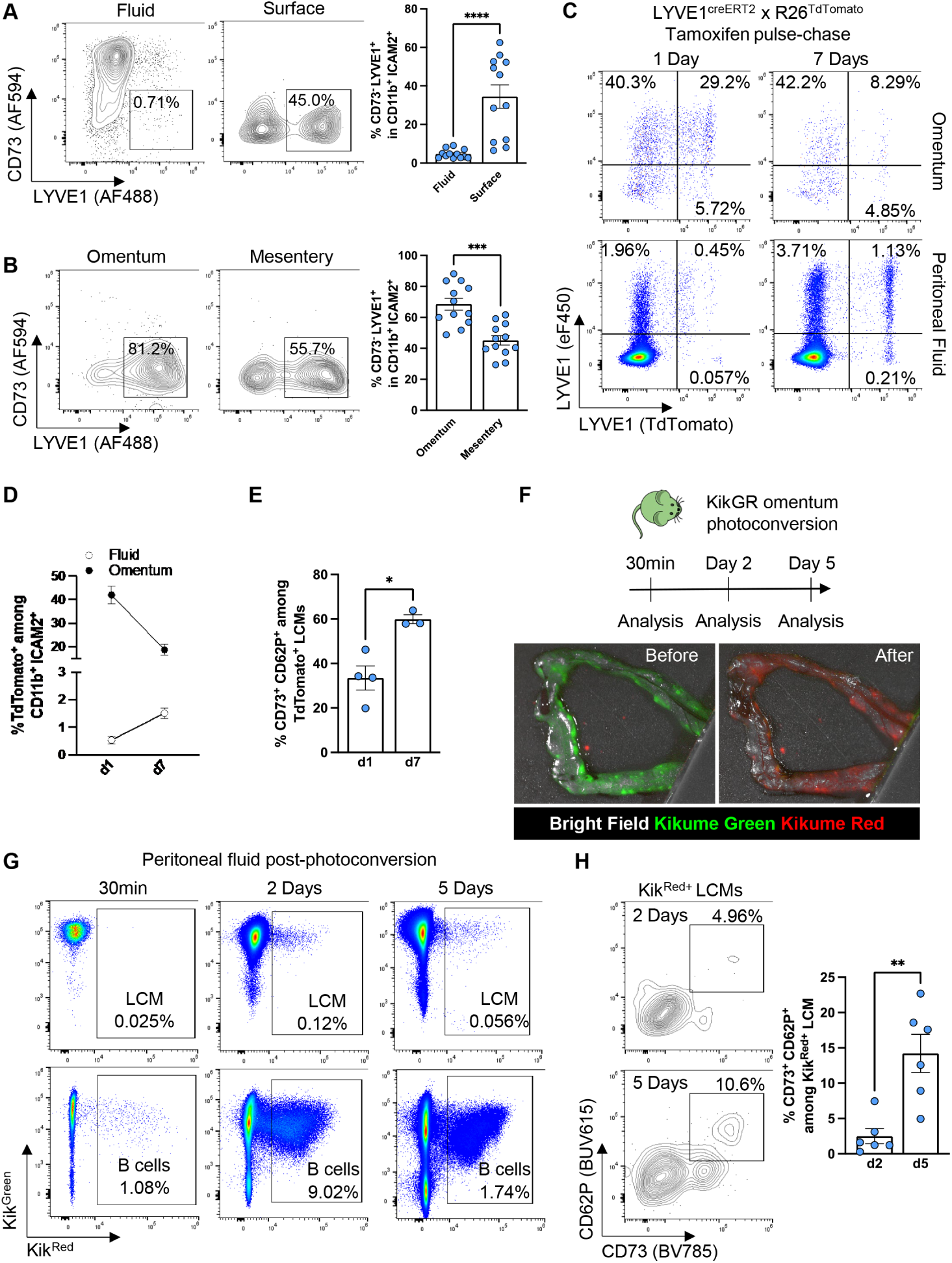
Peritoneal surface-associated ICAM2^+^ peritoneal macrophages contribute to the peritoneal fluid compartment and mature into CD62P^+^ CD73^+^ LCMs. (**A, B**) Representative flow cytometry plots and quantification of CD73^-^ LYVE1^+^ cells among CD11b^+^ ICAM2^+^ LCMs in (A) the peritoneal fluid and mesothelial surfaces or (B) omentum and mesentery. (**C**) Representative flow cytometry plots showing the expression of TdTomato in omental (top row) and peritoneal (bottom row) macrophages from Lyve1^creER^ x R26^TdTomato^ mice at day 1 or day 7 after tamoxifen gavage. (**D**) Proportions of TdTomato^+^ cells among CD11b^+^ ICAM2^+^ macrophages in peritoneal fluid and omentum 1 (n=4) and 7 (n=3) days post tamoxifen gavage. Data were pooled from two independent experiments. (**E**) Proportions of CD62P^+^CD73^+^ macrophages among TdTomato^+^ CD11b^+^ICAM2^+^ macrophages in peritoneal fluid 1 (n=4) and 7 (n=3) days following tamoxifen oral gavage. Data were pooled from two independent experiments. (**F**) Experimental scheme used to photoconvert and chase omental cells for 5 days (top) and representative stereoscope pictures showing the detection of Kikume^Green^ and Kikume^Red^ in the omentum before and after photoconversion (bottom). (**G**) Representative flow cytometry plots showing the detection of Kikume^GREEN^ and Kikume^RED^ in peritoneal macrophages (top row) and B cells (bottom row) over time after photoconversion of the omentum. (**H**) Representative flow cytometry plot (left) and proportions (right) of CD62P^+^ CD73^+^ cells among Kikume^RED+^ peritoneal macrophages 2 (n=6) and 5 (n=6) days after photoconversion. Data were pooled from two independent experiments. Mann-Whitney tests were used for statistical analysis. See also Extended Data Figure 2. *P<0.05, ***P<0.001, ****P<0.0001.

To test whether ICAM2^+^ LYVE1^+^ omental or mesenteric macrophages later migrate into the peritoneal fluid, we performed fate mapping using LYVE1^creER^ R26^TdTomato^ mice. After a single oral tamoxifen administration to pulse-label LYVE1^+^ macrophages, we analyzed the omentum as a representative peritoneal surface and examined the peritoneal lavage fluid over time. Around 35-40% of omental ICAM2^+^ macrophages were TdTomato^+^ 1 day after tamoxifen administration, and this proportion decreased to about 20% 7 days after treatment (**Fig. 2C-D**). In this same interval, the proportion of TdTomato^+^ cells rose in the peritoneal fluid, albeit remaining a small proportion of labeled LCM (**Fig. 2C-D**). The proportion of TdTomato^+^ LCMs co-expressing of CD62P and CD73 increased over time in the peritoneal fluid (**Fig. 2E**). These data are consistent with the concept that LYVE1 expression is transient among some LCMs, particularly those localized to peritoneal surfaces such as the omentum and mesentery, which later mature into CD73^+^CD62P^+^ LCMs within the peritoneal fluid.

To further test whether macrophages that interact with mesothelial surfaces could contribute to the non-adherent ICAM2^+^ macrophage pool in the peritoneal fluid, we used KikGR mice that ubiquitously expressing the Kikume green-red photoconvertible fluorescent protein (*37*) to photoconvert the omentum as a representative peritoneal surface and track photoconverted cells over time (**Fig. 2F**). KikGR mice were anesthetized using isoflurane before carefully exposing the omentum through a 1-cm surgical opening along the linea alba in the peritoneum. The omentum was positioned on a silicon stage, and a second piece of silicon was used to cover the peritoneal opening to block non-targeted photoconversion (**Fig. S2A**). Saline was used to wash away any loosely adherent cells from peritoneal fluid and prevent dehydration of the omentum. Exposure of the omentum to 365 nm light for 3 seconds resulted in instantaneous photoconversion of Kikume from green (Kik^Green^) to red (Kik^Red^) (**Fig. 2F**). The omentum was placed back into the peritoneal cavity, and the peritoneal compartment was resealed with sutures. Mice were then allowed to recover, and the peritoneal lavage was collected and analyzed 30 minutes or 2-5 days later (**Fig. 2F**). Omental CD45^+^ cells displayed Kik^Red^ signal assessed by flow cytometry 30 minutes after photoconversion, while blood monocytes were unlabeled, further confirming the success and specificity of the procedure (**Fig. S2B**). No Kik^Red+^ LCMs were detected in the peritoneal lavage at this early timepoint, but Kik^Red+^ B cells comprised about 1% of B cells in the lavage within 30 minutes (**Fig. 2G**). By 2 days after photoconversion, ICAM2^+^ macrophages in the peritoneal fluid were identified as Kik^Red+^ cells arising from the omentum, comprising about 0.1% of peritoneal fluid LCMs. At day 2, about 10% of peritoneal fluid B cells were Kik^Red+^ (**Fig. 2G**). Kik^Red+^ cells were still present 5 days after the procedure (**Fig. 2G**). Kik^Red^ labeling disappeared from omental ICAM2^+^ macrophages but not from peritoneal lavage LCMs over time (**Fig. S2C**). Time-course analysis revealed that Kik^Red+^ macrophages recovered from peritoneal fluid between days 2 and 5 after photoconversion became progressively more mature, with a higher proportion of CD73^+^CD62P^+^ LCMs at day 5 compared with day 2 (**Fig. 2H**). Thus, these three experimental approaches reveal differentiation of mesothelial surface CD73^-^LYVE1^+^ LCMs into peritoneal fluid CD73^+^CD62P^+^ LCMs.

### Integrin signaling disruption promotes peritoneal fluid accumulation of ICAM2^+^CD73^-^LYVE1^+^ LCMs

Considering the evidence that interactions with mesothelial tissues occur during the differentiation of CD73^-^ LYVE1^+^ LCMs to CD73^+^CD62P^+^ LCMs, we investigated whether manipulations affecting integrin-mediated adhesion would impact this differentiation. Accordingly, we used Lyz2^cre^ x Talin1^fl/fl^ mice (Lyz2^ΔTalin^) to impair inside-out integrin signaling (*1*) as a potential means to diminish macrophage adherence to the omentum and other mesothelial linings. As a positive control, we administered a high dose of zymosan to Lyz2^ΔTalin^ mice and littermate control mice to trigger the macrophage disappearance reaction and recruitment to the omentum. Following this treatment, the number of omental CD11b^+^ICAM2^+^ macrophages was reduced by about half (**fig. S3A**), as expected (*1*). This result confirmed the reduced adherence capacity of LCMs from Lyz2^ΔTalin^ mice.

Under homeostatic conditions and using unsupervised clustering of concatenated data from Lyz2^ΔTalin^ mice versus control littermates (Talin1^fl/fl^ mice without Cre expression) to analyze peritoneal fluid cell phenotypes, it was clear that in the absence of Talin1 expression, CD73^-^ LYVE1^+^ ICAM2^+^ LCMs were much more numerous than in controls (**Fig. 3A**). These cells were CD206^+^, expressed intermediate rather than high levels of F4/80, and expressed high levels of folate receptor β (**Fig. 3B**). Supervised analysis reinforced the increase in this population (**Fig. 3B**). Quantitively, the numbers of CD62P^+^CD73^+^ LCMs (**Fig. 3C**) and peritoneal monocytes (**Fig. S3B**) in male and female Lyz2^ΔTalin^ mice were comparable to those in control littermates, as were the numbers of ICAM2^+^CD73^-^LYVE1^-^ macrophages (**Fig. 3D**). By contrast, the accumulation of ICAM2^+^CD73^-^ LYVE1^+^ macrophages in peritoneal fluid from Lyz2^ΔTalin^ mice was substantial, especially in male mice, in which the numbers of ICAM2^+^CD73^-^LYVE1^+^ macrophages were nearly an order of magnitude higher (**Fig. 3D**), indicating that the CD73^-^LYVE1^+^ mesothelium-associated subset was selectively regulated by loss of the integrin signaling intermediate Talin1. While the numbers of CD73^-^LYVE1^+^ LCMs in peritoneal fluid accumulated when the cells lacked Talin1 expression, the numbers of CD73^-^LYVE1^+^ LCMs in the omentum (**Fig. 3E**) or other peritoneal surfaces (**Fig. 3F**) appeared unaffected.

**Figure 3.**
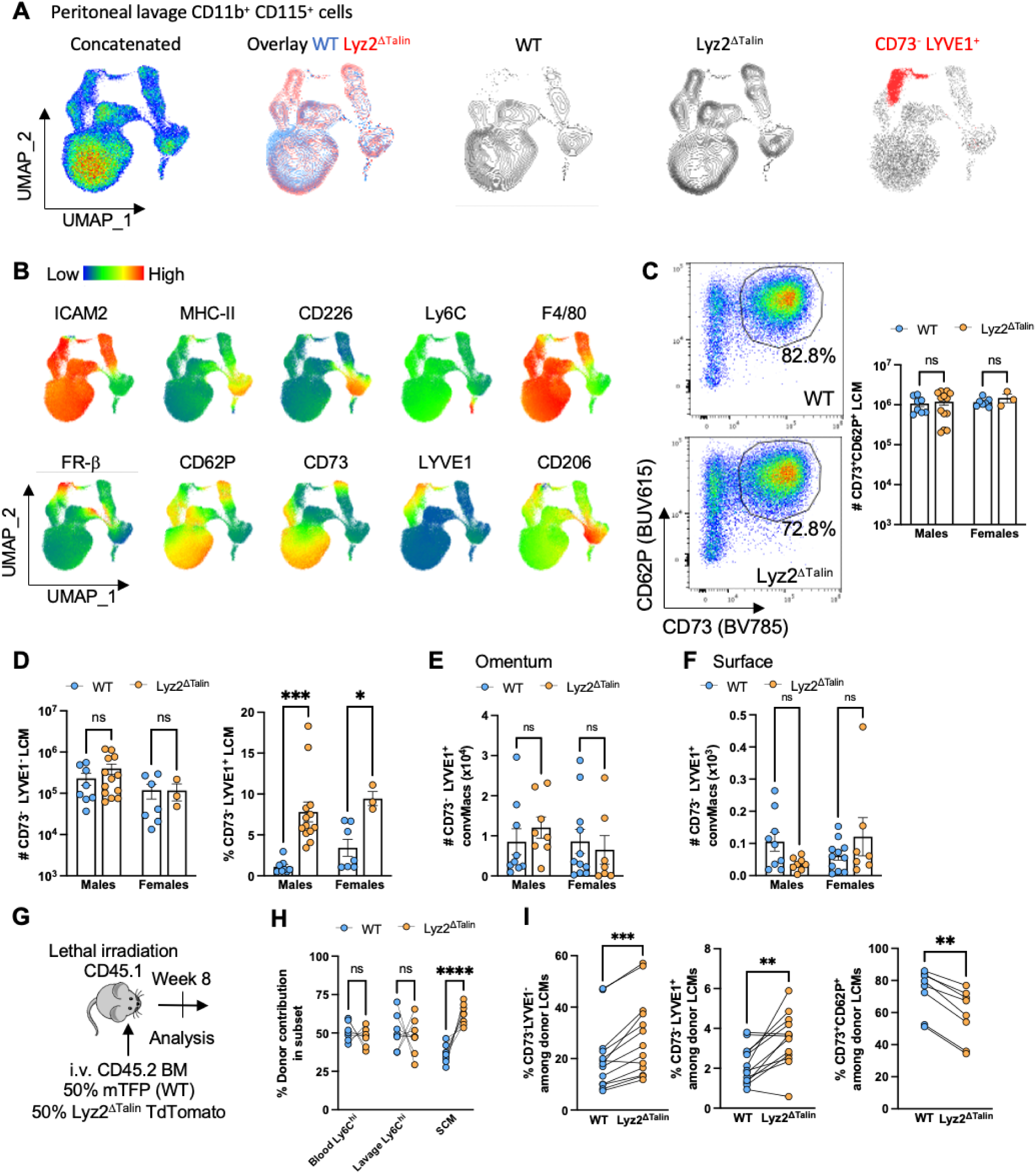
Integrin signaling disruption promotes peritoneal fluid accumulation of ICAM2^+^CD73^-^LYVE1^+^ LCMs. (**A**) UMAP projection of peritoneal CD11b^+^ CD115^+^ cells concatenated from WT (Lyz2^+/+^ Talin1^fl/fl^) and Lyz2^ΔTalin^ (Lyz2^cre/+^ Talin1^fl/fl^) mice. (**B**) UMAP representation of surface marker expression in the concatenated dataset from (A). (**C, D**) Representative flow cytometry plots and quantification CD73^-^ LYVE1^+^ (C), CD73^-^ LYVE1^-^ and CD62P^+^ CD73^+^ (D) macrophages in peritoneal fluid from WT (male n=14, female n=7) and Lyz2^ΔTalin^ (male n=24, female n=3) mice. Data were pooled from three independent experiments. (**E**, **F**) Quantification of CD73^-^ LYVE1^+^ LCMs, previously described as converting macrophages (convMacs), in the omentum (E) and mesothelial surfaces (F) from WT (male n=9, female n=8) and Lyz2^ΔTalin^ (male n=8, female n=7) mice. (**G**) Experimental scheme used for competitive transplant of WT (Actin^mTFP^) and Lyz2^ΔTalin^ bone marrow into lethally irradiated CD45.1 recipients. (**H, I**) Proportions of WT and Lyz2^ΔTalin^ donor-derived cells in monocytes and SCMs (H), and CD11b^+^ICAM2^+^ LCM subsets (I). Data from recipient mice (n=8) were analyzed and combined across two independent experiments. ANOVA (panels C, D, E, F, and H) and Mann-Whitney (panel I) tests were used for statistical analysis. See also Extended Data Figure 3. *P<0.05, **P<0.01,***P<0.001.

Observing that the increased accumulation of Talin1-deficient LYVE1^+^ LCMs in the peritoneal fluid did not increase the number of mature CD73^+^CD62P^+^ LCMs retrieved from the peritoneal cavity as we expected, we wondered if LYVE1^+^ LCMs and CD73^+^CD62P^+^ LCMs may not follow a strict precursor-product relationship as earlier envisioned. We also considered that the presence of embryonically derived CD73^+^CD62P^+^ LCMs might obscure the evaluation of the relationship between the two phenotypes. Thus, to eliminate concerns about the presence of embryonically derived LCMs, we set up a competitive BMT wherein lethally irradiated host mice received a 1:1 mixture of WT bone marrow cells marked with a universally expressed teal fluorescent protein (mTFP1) and Lyz2^cre^ xTalin^fl/fl^ xR26^TdTomato^ bone marrow cells (**Fig. 3G**). As a control, leukocytes in the blood compartment, including Ly6C^hi^ monocytes (**Fig. 3H**), were equally represented in a 1:1 ratio and maintained this 1:1 ratio in the peritoneal lavage (**Fig. 3H**). Talin1 deficiency supported a shift toward a higher proportion of SCMs, suggesting that, in the absence of Talin1, some aspect of differentiation beyond monocyte recruitment was favored for this macrophage subset (**Fig. 3H**). When we plotted the ratios of LCM differentiation intermediates, including CD73^-^LYVE1^-^ or CD73^-^LYVE1^+^ cells in the LCM differentiation pathway, Talin1 deficiency favored accumulation of peritoneal fluid CD73^-^LYVE1^+^ LCMs (**Fig. 3I**). By contrast, WT cells were conversely dominant among the most mature LCMs co-expressing CD73 and CD62P (**Fig. 3I**). Overall, data arising from BMT (**Fig. 3 G-I**) generated similar conclusions as in the steady state (**Fig. 3A-F**), showing that Talin1 deficiency impacted the distribution of CD73^-^LYVE1^+^ LCMs in the peritoneum, while having little or even the opposite impact on the accumulation of CD73^+^CD62P^+^ LCMs in peritoneal fluid.

### LYVE1^+^ LCMs are minor precursors of CD73^+^CD62P^+^ LCMs even after bone marrow transplant

The findings above motivated us to formally evaluate the presumed relationship between CD73^-^LYVE1^+^ LCMs and CD73^+^CD62P^+^ LCMs. In particular, we wondered if a LYVE1^+^ stage was required to generate CD73^+^CD62P^+^ LCMs from bone marrow cells. To evaluate the proportion of LCMs arising from bone marrow-derived LYVE1^+^ progenitors in the context of BMT, CD45.1 C57BL/6 congenic mice were lethally irradiated and transplanted with YFP^-^ bone marrow from LYVE1^cre^ x R26^YFP^ mice (**Fig. 4A**). YFP^-^ LYVE1^cre^ x R26^YFP^ mice bone marrow donors were either Gata6^fl/+^ or Gata6^fl/fl^, enabling a detailed examination of the relationship between LYVE1 and Gata6 in the differentiation of various stages of ICAM2^+^ LCMs (**Fig. 4A**). At 8 weeks following BMT, we gated CD73^-^ LYVE1^+^ or CD73^-^LYVE1^-^ LCMs or CD73^+^CD62P^+^ mature LCMs. Macrophages with *Gata6* deleted had reduced numbers of CD73^+^CD62P^+^ LCMs, as expected (*15, 16*) (**Fig. 4B**), while both CD73^-^LYVE1^-^ and CD73^-^LYVE1^+^ LCMs increased in number in the absence of *Gata6* (**Fig. 4B**). The percentage of LYVE1^+^ LCMs that expressed YFP was greater than 75% (**Fig. 4C**), but despite that, only about 10% of CD73^+^CD62P^+^ LCMs were YFP^+^ (**Fig. 4C**), and, in even in *Gata6^+^* mice, YFP^-^CD73^+^CD62P^+^ LCMs outnumbered by an order of magnitude all other stages of LCM differentiation as well as all LCMs bearing the YFP^+^ lineage tracer (**Fig. 4D**). These data indicate that, unexpectedly, only a few CD73^+^CD62P^+^ LCMs arise from CD73^-^LYVE1^+^ LCMs. Furthermore, although it was evident that the LYVE1^+^ ‘converting LCM’ certainly gives rise to a mature LCM, that few mature LCMs rely on a LYVE1^+^ intermediate upends the notion that a LYVE1^+^ progenitor is the sole or main progenitor of mature LCMs.

**Figure 4.**
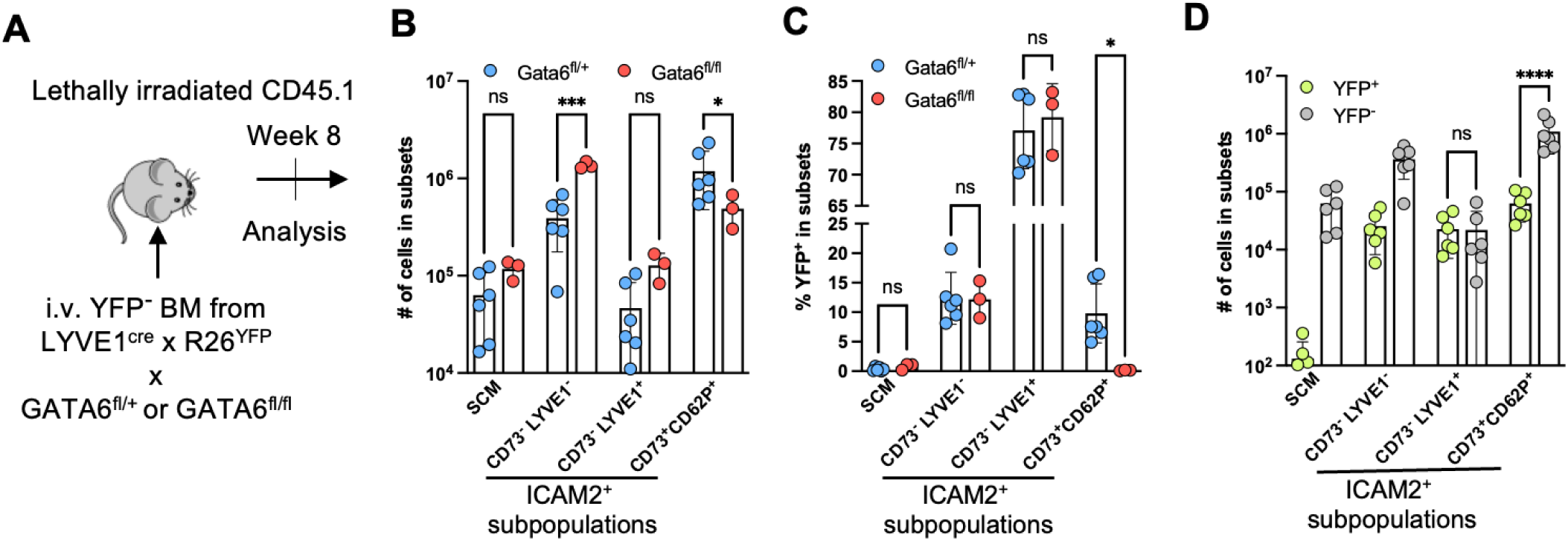
LYVE1^+^ LCMs are minor precursors of CD73^+^CD62P^+^ LCMs even after bone marrow transplant. (**A**) Experimental scheme used to generate BMTs for fate mapping cells that express LYVE1 during differentiation after transplant. (**B, C**) (B) Quantification of donor SCMs and ICAM2^+^ LCM subsets and (C) YFP expression from CD45.1 mice transplanted with YFP^-^ bone marrow cells from Lyve1^cre^ x R26^YFP^ mice with conditional deletion of Gata6 (Gata6^fl/fl^, n=3) or control littermates (Gata6^fl/+^, n=7). (**D**) Quantification of donor YFP^+^ and YFP^-^ cells among SCMs and ICAM2^+^ LCM subsets from CD45.1 mice transplanted with YFP^-^ bone marrow cells from Lyve1^cre^ x R26^YFP^ mice. Each symbol represents data from an independent mouse. An ANOVA was used for statistical analysis. (E) Quantification of CD62P+CD73+ cells in the peritoneal cavity of mice administrated with low dose (10ug) zymosan with (OMX) or without (Sham) omentectomy. *P<0.05, **P<0.01, ***P<0.001, ****P<0.0001, n.s. not significant.

### Bone marrow-reconstituted CD73^-^ but not CD73^+^ LCMs quantitatively depend on monocytes

LYVE1^+^ LCMs appear to arise from monocytes, especially following the onset of inflammatory conditions in serosal cavities (*18*). Having unexpectedly gathered evidence that, even after BMT, LYVE1^+^ LCMs were not obligate precursors for CD73^+^CD62P^+^ LCMs, we wondered if the concept and evidence suggesting that monocytes were the precursors for all LCMs after BMT (*25*) warranted more detailed exploration. Furthermore, we wondered if, in the steady state wherein embryonically derived LCMs can proliferate for self-sustenance (*7*), the LCM compartment would show alterations in mice with greatly reduced blood monocytes.

To address the latter point first, we analyzed mice lacking blood monocytes due to the introduction of three point mutations in NFIL3 binding sites within the Zeb2 gene locus (*33*). We refer to this strain herein as triple-mutant of Zeb2 (Zeb2^TM^) mice. After confirming the loss of blood monocytes (**Fig. S4A**) and applying the gating strategy shown in **Fig. S1A**, newly recruited monocytes within the peritoneum (**Fig. 5A**), and the CD226^+^ SCMs (**Fig. 5B**) previously described to arise from monocytes (*10*) were greatly reduced in resting Zeb2^TM^ mice (**Fig. 5A-B**). When focusing on ICAM2^+^ LCMs, we observed no reduction in CD73^+^CD62P^+^ LCM in Zeb2^TM^ mice (**Fig. 5C**). However, there was a marked reduction in all ICAM2^+^ CD73^-^ LCMs in Zeb2^TM^ mice (**Fig. 5C**). The subpopulation of CD73^-^LYVE1^+^ LCMs was likewise substantially reduced (**Fig. 5C**). Failure to observe reduced CD73^+^CD62P^+^ macrophages might be expected in homeostasis, since LCM self-renew after being seeded by embryonic progenitors before birth (*7*) (*10*). Nonetheless, these data indicated that there is a low background of monocyte recruitment to the peritoneal cavity during homeostasis, giving rise to CD73^-^LYVE1^-^ICAM2^+^ LCMs and CD73^-^ LYVE1^+^ICAM2^+^ LCMs without impacting CD73^+^CD62P^+^ LCMs. We enumerated the different subpopulations of LCMs after induction of peritoneal inflammation with low dose zymosan to increase monocyte recruitment (*11*). This stimulus elevated the numbers of the CD73^-^LYVE1^+^ LCMs 2 weeks later while not impacting the overall numbers of CD62P^+^CD73^+^ LCMs (**Fig. S4B**). This finding supports the concept that CD73^-^LYVE1^+^ cells likely originate from recruited monocytes and aligns with the lack of correlation between the numbers of CD73^-^LYVE1^+^ LCMs and CD73^+^CD62P^+^ LCMs apparent in our studies of Talin1-deficient macrophages (**Fig. 4**).

**Figure 5.**
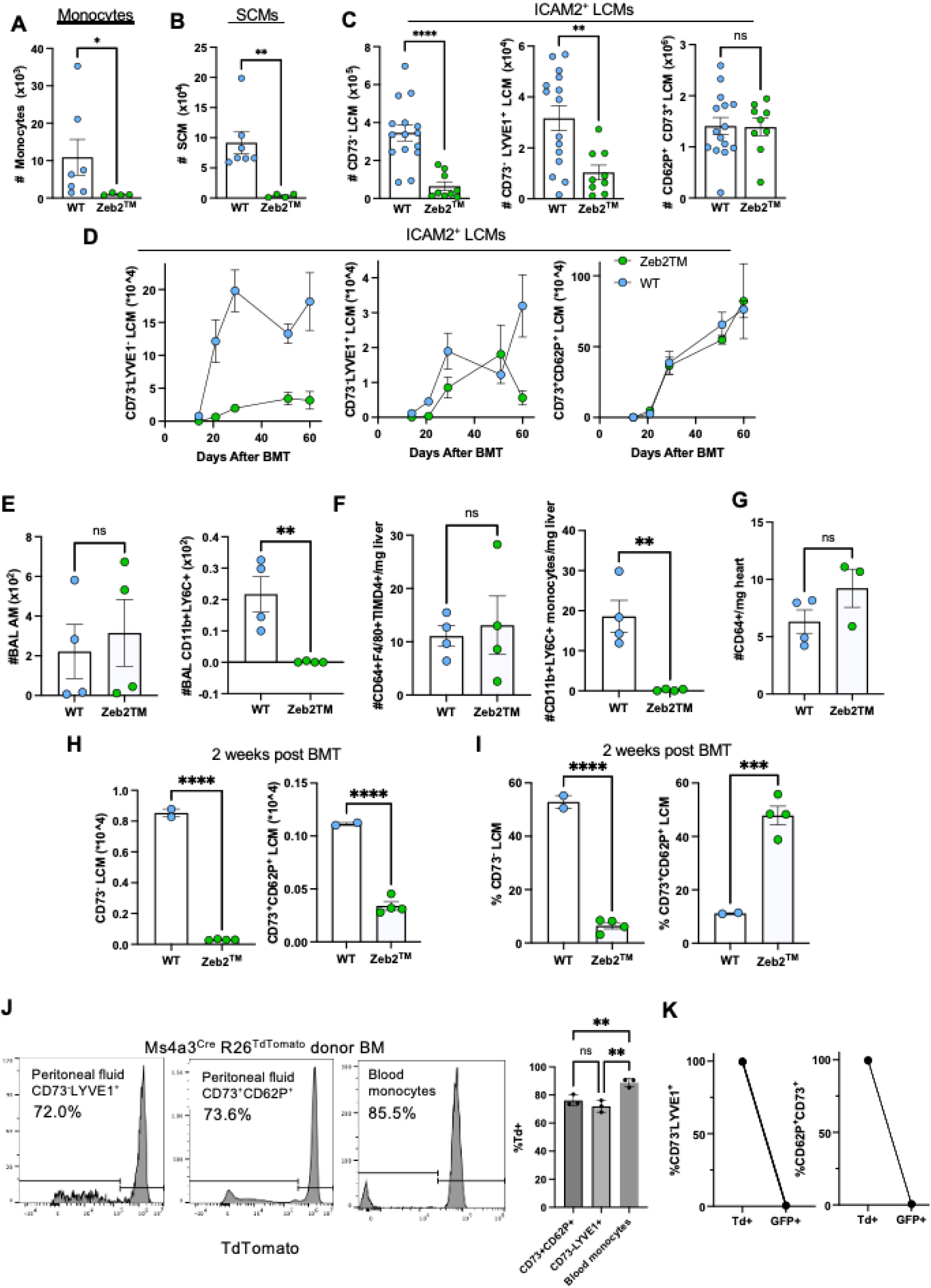
Reconstitution of different peritoneal macrophage subsets across time after BMT. (A-B) Quantification of peritoneal (A) monocytes, and (B) SCMs in WT control (n=7) and Zeb2^TM^ (n=4) mice at steady state. Data are representative of 3 independent experiments. **(C)** Quantification of total CD73^-^ LCM (left), CD73^-^ LYVE1^+^ LCM (middle), and CD62P^+^ CD73^+^ (right) LCMs in WT control (n=15) and Zeb2^TM^ (n=9) mice at steady state. Data were pooled from two independent experiments. Mann-Whitney tests were used for statistical analysis. **(D)** Quantification of CD73^-^LYVE1^-^ (left), CD73^-^LYVE1^+^ (middle) and CD62P+ CD73+ (right) LCMs at 14 days (WT n=2, Zeb2^TM^ n=4), 21 days (WT n=3, Zeb2^TM^ n=4), 29 days (WT n=3, Zeb2^TM^ n=4), 51 days (WT n=4, Zeb2^TM^ n=3), and 60 days (WT n=9, Zeb2^TM^ n=6) post BMT. WT group (blue dots) are CD45.1 recipient mice transplanted with 50:50 Lyz2^cre^ x R26^TdTomato^ and CCR2^gfp/gfp^ bone marrow after lethal irradiation. Zeb2^TM^ group (green dots) are CD45.1 recipient mice transplanted with Zeb2^TM^ bone marrow after lethal irradiation. (**E-G**) Quantification of (E) Bronchoalveolar lavage alveolar macrophages (BAL AM) and monocytes, (F) liver Kupffer cells and monocytes, and (G) heart macrophages 4 weeks after transplantation of lethally irradiated CD45.1 mice with WT or Zeb2TM bone marrow. **(H-I)** The cell number quantification (H) and percentage of total ICAM2+ LCMs (I) of the CD73^-^ LCMs and CD62P^+^ CD73^+^ LCMs in the peritoneal cavity of lethally irradiated CD45.1 recipient mice 21 days post transplantation with bone marrow of different genotypes. Multiple two-tailed t-tests followed by FDR correction; **P < 0.01, ***P < 0.001, ****P < 0.0001, ns. not significant. **(J)** Representative plots and bar plot showing the proportions of TdTomato^+^ cells among CD73- LYVE1+, CD73+CD62P+ LCMs or blood monocytes in mice transplanted with Ms4a3^Cre^R26^TdTamato^ donor BM. **(K)** Proportions of TdTomato^+^ versus GFP^+^ cells out of the total CD73^-^LYVE1^+^ or CD62P^+^CD73P^+^ LCMs in CD45.1 recipient mice transplanted with 50:50 Lyz2^cre^ x R26^TdTomato^ and CCR2^gfp/gfp^ bone marrow after lethal irradiation. *P<0.05, **P<0.01, ***P<0.001, ****P<0.0001.

To eliminate the complexity of embryonically seeded LCMs as a variable, we next employed a model of genotoxic stress induced by whole-body irradiation and BMT, which results in the complete replacement of LCMs by donor-derived bone marrow cells(*7*), as shown in Figure 4. Bone marrow from Zeb2^TM^ (CD45.2^+^), WT mice, or Lyz2^cre^ x R26^TdTomato^ reporter mice was transplanted into congenic CD45.1**^+^** C57BL/6 host mice for comparison. Tracking the congenic CD45 marker allowed for the analysis to be exclusively focused on donor-derived macrophages. Because replenishment of LCMs occurs over several weeks, we harvested mice at the earliest time when replenishment is just starting, 2 weeks after lethal irradiation (*7*), and at various time points thereafter up to 8 weeks (**Fig. 5D**). As a positive control, mice receiving Zeb2^TM^ bone marrow had many fewer recruited monocytes in the peritoneal cavity compared with mice receiving WT donor cells, albeit the harvest at 6 weeks revealed a relative uptick in peritoneum-entering monocytes from Zeb2^TM^ donors compared to other time points (**Fig. S4C**). Zeb2^TM^ donor marrow also resulted in the expected loss of SCMs (*10*) in the peritoneum compared with mice receiving control bone marrow, confirming the quantitative reliance of SCMs on monocytes (**Fig. S4D**). CD73^-^ LCMs were greatly diminished in mice receiving Zeb2^TM^ donor marrow compared with those receiving WT donor cells (**Fig. 5D**). CD73^-^LYVE1^+^ LCMs showed an uptick at 6 weeks with a drop again at 8 weeks (**Fig. 5D**) that paralleled the monocyte uptick in the cavity (**Fig. S4C**), altogether suggesting that CD73^-^ LCMs were collectively closely reliant on the quantitative supply of monocytes for their replenishment. In stark contrast, the repopulation of CD73^+^CD62P^+^ mature LCMs was largely intact in mice reconstituted with Zeb2^TM^ donor marrow (**Fig. 5D**). Counterintuitively, while the most mature LCMs were scarcely impacted by the diminished pool of monocytes in Zeb2^TM^ BMT mice, the presumed precursors of these LCMs were markedly hobbled. The quantitative lack of reliance on the availability of recruited monocytes was also evident in mice receiving Zeb2^TM^ donor marrow when we examined alveolar macrophages, Kupffer cells, and cardiac macrophages, as all these resident macrophages were replenished from Zeb2^TM^ marrow donors as efficiently as from WT marrow donors 4 weeks after BMT, even though monocytes were not detected in the same organs when Zeb2^TM^ donor marrow was supplied (**Fig. 5E-G**)

Since donor replenishment of peritoneal macrophages is just getting underway at 2 weeks post-irradiation (**Fig. 5D**), this time point serves as an early window into the different stages of macrophage differentiation. As the data from this timepoint were quantitatively buried under the greater replenishment of LCMs that followed in the later weeks, we replotted the 2-week timepoint in separate bar graphs. **Fig. 5H-I** depict the repopulation of CD73^-^ LCMs and CD73^+^CD62P^+^ LCMs at the 2-week timepoint. At this early stage, the total number of all LCMs, expressing CD73 or not, was reduced in Zeb2^TM^ mice (**Fig. 5H**). However, the severity of the reduction relative to controls was less for CD73^+^CD62P^+^ LCMs than for CD73^-^ LCMs (**Fig. 5H-I**). Of the LCMs recovered at 2 weeks, more than 40% were CD73^+^CD62P^+^ in the Zeb2^TM^ mice. In contrast, only about 10% of the LCMs at 2 weeks had matured to CD73^+^CD62P^+^ LCMs in control mice (Lyz2^cre^ x R26^TdTomato^) with normal numbers of monocytes (**Fig. 5I**). To formally test whether these LCM populations arose from granulocyte-macrophage progenitors (GMP) after BMT, we reconstituted lethally irradiated mice with Ms4a3^cre^ x R26^TdTomato^ bone marrow (**Fig. 5J**). When we examined whether CD73^+^CD62P^+^ LCMs were TdTomato^+^ in this experiment, we found that the fraction of TdTomato^+^ CD73^+^CD62P^+^ LCMs was about 70%, resembling CD73^-^LYVE1^+^ LCMs, while blood monocytes were labeled at around 85% (**Fig. 5J**). Furthermore, both CD73^+^CD62P^+^ and CD73^-^LYVE1^+^ LCMs were reliant on CCR2^+^ cells for replenishment since in a BMT setting wherein Lyz2^cre^ x R26^TdTomato^ and CCR2^gfp/gfp^ knock-in/knockout marrow was given 1:1 resulted in strong skewing toward Td^+^ (>99%) over GFP^+^ LCMs regardless of whether CD73^-^ or CD73^+^ LCMs were examined (**Fig. 5K**). These data collectively demonstrate that CD73^+^CD62P^+^ LCMs arise mainly from a GMP-derived CCR2-dependent myeloid progenitor, possibly a monocyte, following irradiation injury. Whether other monocyte lineages such as those arising from MDPs are responsible for the generation of TdTomato^-^ cells remains to be tested. However, if monocyte recruitment to the peritoneal cavity is, in fact, necessary in the initial stage of CD73^+^ CD62P^+^ LCM replenishment, this population requires minimal progenitor influx. On the contrary, the pools of SCMs and CD73^-^ LCMs, including CD73^-^LYVE1^+^ LCMs, sustain only proportionally to the extent of monocyte influx, indicating that these LCMs never acquire tissue residency. Thus, CD73^-^LYVE1^+^ LCMs and CD73^+^CD62P^+^ LCMs appear to exist largely as distinct pools with some overlap based on evidence from the LYVE1-based lineage tracing that shows CD73^-^LYVE1^+^ LCMs support the development of a minority of CD73^+^CD62P^+^ LCMs.

## DISCUSSION

Past literature supported the notion that monocytes can migrate into the peritoneal cavity, where they differentiate into mature LCMs that closely resemble LCMs derived from embryonic progenitors (*25, 26*). Differentiation intermediates that repopulate LCMs at first express intermediate levels of F4/80, contrasting with the high levels on the mature Gata6^+^ LCM (*11, 16, 19, 25, 29*). These intermediates upregulate FRβ, CD206, and LYVE1 (*16, 18*) before they induce high Gata6, CD73, and CD62P levels. At this point, they downregulate LYVE1, CD206, and FRβ, such that the latter group of genes is associated with the transient period in their conversion from monocytes to mature LCMs (*18*). In this study, we set out to define how these differentiation steps relate to the interaction of monocyte-derived cells with the mesothelial lining of the peritoneal cavity, including specialized peritoneal folds like the omentum. A previous study identified LCM differentiation intermediates accumulating in the omentum when LCM terminal differentiation was blocked by genetic deletion of Gata6 (*15*). Employing a mouse model wherein integrin binding to ligands could be broadly blocked by macrophage-selective deletion of the inside-out signaling adaptor Talin1 (*38*), along with short-term fate-mapping through photoconversion and inducible tagging of LYVE1^+^ expressing macrophages, our findings support the framework that mesothelial tissues of the peritoneal cavity facilitate key steps in monocyte differentiation to LCMs. We show that the LYVE1^+^ LCM, previously termed the ‘converting LCM’, (*18, 31*) mobilizes from tissues covered in mesothelium like the omentum to the peritoneal fluid, where it can differentiate further into CD73^+^CD62P^+^ mature LCMs that coincide with Gata6^+^ and Gata6-dependent LCMs. These findings align well with established paradigms in the field regarding the developmental support of LCMs from mesothelial surfaces (*12, 23, 24*) and the role of LYVE1^+^ LCMs as precursors to Gata6^+^CD73^+^CD62P^+^ mature LCMs (*18*). However, our findings also challenge the idea that the maturation sequence of CD73^-^LYVE1^+^ LCMs into CD73^+^CD62P^+^ LCMs involving integrin-mediated adhesion to peritoneal surfaces like the omentum followed by a transition into peritoneal fluid, respectively, is the singular mechanism required to generate the majority of CD73^+^CD62P^+^ LCMs, even after enforced replacement of LCMs through BMT.

When we observed that the loss of Talin in LCMs greatly impacted LYVE1^+^ LCMs but not mature LCMs, we carried out fate mapping studies that surprisingly revealed that LYVE1^+^ LCMs were minor intermediates in repopulating mature LCMs, not major intermediates as they previously seemed (*18*). Experiments to probe the role of monocytes in repopulating LCMs recapitulated the literature that mature LCM repopulation relies on CCR2 (*25*) and is largely linked to precursors that develop through a stage involving the expression of Ms4a3 (*26*). However, approaches that crippled the development of monocytes per se due to mutations in the Zeb2 enhancer (*32, 33*) uncovered a marked discrepancy between the accumulation of LCM differentiation intermediates, which were minimal in the Zeb2 mutant lines, and the repopulation of mature LCMs that were only modestly impacted in the Zeb2 mutant lines. These data quantitatively uncouple LCM differentiation intermediates from the repopulation of mature LCMs, revealing that a single linear pathway in a precursor-product relationship from monocytes to mature LCMs does not exist. Instead, we argue for two possibly interconnected pathways in the bone marrow-dependent repopulation of LCMs from monocytes. First, a pathway of LCM differentiation exists that is entirely reliant on monocytes. This pathway includes the differentiation of monocytes to ICAM2^+^LYVE1^+^CD73^-^ intermediates, and these can give rise to at least some Gata6^+^ LCMs. However, a second pathway does not give rise to the LYVE1^+^ differentiation intermediate but ultimately gives rise to a vastly greater number of Gata6^+^ LCMs than the first pathway. This second pathway does depend on CCR2 and involves the recruitment of a myeloid progenitor that has earlier passed through the Ms4a3^+^ GMP bone marrow progenitor stage (*26*). It is possible, but not entirely certain, that this second pathway arises from monocytes, like the first pathway. If it does, we argue that the recruited monocytic precursors deviate from quantitative reliance on monocytes to give rise to LCMs. One way in which this might happen is if a particular pathway of differentiation allowed monocytes to massively proliferate before expressing LYVE1, similar to the phenomenon that was recently observed in lungs (*39*), repopulating LCMs in a manner not reliant on the same Zeb2 enhancer as the LCM that arises in the first pathway. Indeed, the acquisition of the program that sustains embryonic macrophages could explain why the second pool of bone marrow-derived LCMs do not depend on Zeb2, since embryonic resident macrophages are sustained in mice bearing Zeb2 enhancer mutations that cripple monocyte and dendritic cells (*32*). Our less extensive examination of macrophages in other tissues like the alveolar compartment, the heart, and the liver suggests that the finding that monocytes are not quantitatively required to replenish tissue-resident macrophages in the peritoneum after BMT applies to other organs. Perhaps rather than the embryonic versus adult bone marrow origin providing the distinguishing feature of Zeb2 dependency, different maturation paradigms may determine Zeb2 dependency. For example, mature intestinal macrophages, which are substantially seeded even in homeostasis by monocytes unlike many resident tissue macrophages (*40–42*), fail to exhibit dependence on Zeb2 (*33*), although a close look at the data suggest that the earlier defined intermediates of intestinal macrophage differentiation (*43*) are reduced (*33*). Additional complexity in macrophage repopulation has recently been linked to whether monocytes arise from GMP or MDP progenitors (*44*). Yet, our finding that most LCMs appear to be linked to a GMP background lineage based on use of the Ms4a3^Cre^ reporter line seems to preclude the possibility that the two pathways we uncover are simply a result of whether the monocyte arises from a GMP progenitor or not. However, this topic deserves more attention in the future. It is worth noting that Geissmann and colleagues pointed to all homeostatic tissues having both embryonically derived and monocyte-derived macrophages within them (*45*). Our findings, together with other work following macrophage intermediates separately from mature macrophages (*43*), raise the possibility many tissue macrophages exist in a duality wherein a shadow of the main self-sustaining population, often of embryonic origin, is secondarily maintained by monocyte-derived intermediates that develop into some of the same mature macrophage endpoints but do not take up residency and may not progress along the exact same lineup of intermediates.

Future studies will be needed to further delineate the steps and requirements of the two pathways described above. We argue that doing so is critical, as it seems likely to us that only the first pathway is operative in humans. That is, the human peritoneal cavity hosts an ongoing recruitment of monocytes and differentiation of LCM intermediates like the LYVE1^+^ LCM (*31*) that we show here is monocyte-dependent in mice, along with a small proportion of Gata6^+^ LCMs (*31*). We were puzzled why the proportion of Gata6+ LCMs was so low in humans (<5%), while it was so high in mice (*31*). Our data raise an explanation that we had not earlier considered. With two pathways operative in mice to replenish Gata6^+^ LCMs, with the dominant pathway being the pathway least reliant on quantitative replenishment from monocytes, it seems reasonable to consider that humans may lack this second pathway that gives rise to LCMs in mice while possessing the first pathway that gives rise to a more modest yield and proportion of Gata6^+^ LCMs. Human LCMs, even those expressing Gata6, do not express CD73 (*31*), fitting with the hypothesis that in both mice and humans, differentiation of CD73^-^ LCMs defines a pathway for generating LCMs that depends on monocytes and gives rise to a modest yield of Gata6^+^ LCM. It is interesting to note that loss of RARγ or RXRα in mouse LCMs leads to an enrichment of LYVE1^+^ LCMs with many fewer remaining Gata6^+^ LCMs in mice. These nuclear receptors are scarcely expressed in human LCM (*31*), so these retinoic acid-linked pathways deserve attention as possible pathways that are highly operative in mice and distinguish LCMs in mice from those in humans. Given the evidence of these two pathways and our collective findings, it makes sense that only the LYVE1^+^ LCM pathway that generates Gata6^+^CD73^+^CD62P^+^ LCMs may rely on mesothelial interactions. That is, our findings support the earlier proposed concept that mesothelial surfaces promote macrophage development (*15*), while they also reveal that the main LCMs in mice do not rely on such interactions. Instead, in the absence of Gata6, the one remaining pathway of macrophage differentiation, enriched in ICAM2^+^Gata6^-^ LCMs, was simply more prominent.

In summary, these studies establish a new model for the repopulation of tissue macrophages by monocytes. Differentiation bifurcates into at least two pathways that support tissue resident mature cells, one more reliant on the quantitative recruitment of monocytes than the other. We have mainly focused on this finding in the peritoneal cavity, where differentiation intermediates to replenish LCMs from monocytes have been characterized. Many details remain unknown, particularly regarding the pathway through which resident macrophages arise from bone marrow progenitors independently of Zeb2, which will be the focus of future work.

## ACKNOWLEDGEMENTS

We are indebted to the expert mouse typing and care provided by Mary Wohltmann and Galina Fedotova. AG is now affiliated with Université Côte d’Azur (France). TL is now affiliated with The Scripps Research Institute (California).

## Author contributions

AG, JH, RLM, CGH, MMC, TTH, XL conducted experiments and analyzed data; AG, JH, RLM, JC, CGH, GJR designed experiments; SD, JK, TL and KMM established key resources; GJR, KMM, KJL, JDS, SCM, and BHZ provided supervision and training, regulatory compliance, and funding; AG, JH, RLM and GJR wrote the manuscript with editing input from all authors.

## MATERIALS and METHODS

### Mice

All C57BL/6J and CD45.1 wild-type (WT) mice were purchased from The Jackson Laboratory. Gata6^fl/fl^ mice (Gata6^tm2.1Sad^/J, Jax 008196) were backcrossed to C57BL6/J background and crossed to Lyve1^cre^ mice (B6;129P2-Lyve1^tm1.1(EGFP/cre)Cys^/J, Jax 012601) or Lyz2^Cre^ (B6.129P2-Lyz2^tm1(cre)Ifo^/J, Jax 004781) and R26^LSL-YFP^ (B6.129X1-Gt(ROSA)26Sor^tm1(EYFP)Cos^/J, Jax 006148) mice. Ms4a3cre mice were purchased from The Jackson Laboratory (C57BL/6J-Ms4a3em2(cre)Fgnx/J, Jax 036382) and crossed to R26^LSL-tdTomato^ mice. Tln1^fl/fl^ mice (*9*), KikGR (Tg(CAG-KikGR)33Hadj/J, Jax 013753) (*37*), and Actin^mTFP^ mice (*46*) were bred and maintained at Washington University. Zeb2^TM^ mice were generated at Washington University and crossed as previously described (*33*). Lyve1^creER^ mice were generated at Washington University as described (*47*). Lyz2^Cre^ (B6;129S-Gt(ROSA)26Sor^tm1(CAG-COX8A/Dendra2)Dcc^/J, Jax 018385) mice (*48*) were also bred to Tln1^fl/fl^ and R26^LSL-tdTomato^ mice. In all experiments involving Cre-mediated deletion of target genes, breeding was done with one parent carrying a single copy of Cre, such that Cre^+^ mice were compared to Cre^-^ littermate controls. Mice were kept in specific pathogen-free conditions maintained by the Washington University School of Medicine Division of Comparative Medicine. Facilities were maintained at an ambient temperature of 23-24°C with a 12/12 light/dark cycle. Mice had access to food and water *ad libitum*. All experiments and procedures were approved by Washington University School of Medicine Institutional Animal Care and Use Committee (protocol # 22-0433).

### Omentectomy

Mice were anesthetized using 1-2% isoflurane with a flow rate of 0.8-1.0 liter/min and placed on a heating pad. The peritoneal cavity was opened along the linea alba, the midline of the peritoneal wall, to avoid bleeding. The greater omentum was exposed and cauterized through this incision.The peritoneal incision was closed with 4-0 Vicryl absorbable continuous sutures and the skin with non-absorbable 5-0 nylon simple interrupted sutures. After administration of a subcutaneous injection of saline and the analgesic buprenorphine, mice were placed in an infant incubator overnight and monitored until recovery.

### Photoconversion

Omental photoconversion was accomplished under general anesthesia by continuous inhalation of 1–2% isoflurane with a flow rate of 0.8-1.0 liter/min. Mice were placed on a heating pad in a supine position. The omentum was carefully exposed through a midline incision in the skin and the linea alba along the peritoneum (see Supp. Figure 3a). Saline was applied to rinse the omentum and keep it hydrated. Photoconversion was achieved by shining a 405nm hand-held laser, positioned at a distance of roughly 5 cm directly above with light targeted to the omentum for 3 seconds. The midline incision was then closed with absorbable continuous sutures in the peritoneum (4-0 Vicryl) and non-absorbable simple interrupted sutures in the skin (5-0 nylon). Mice were resuscitated with a subcutaneous injection of saline and buprenorphine analgesia. They were allowed to recover in a warmed incubator.

### Bone marrow transplant (BMT)

Recipient mice were lethally irradiated with 800 cGy in an x-ray irradiator with a turn table to distribute the irradiation across mice. Forelimbs and hindlimbs from donor mice were isolated, and bone marrow was collected in a class II laminar flow hood to maintain sterility. The long bones were mechanically separated from the muscles and placed in RPMI-1640 with 5% FBS to maintain cell viability. They were then soaked in 70% ethanol for sterilization and returned to RPMI-1640 with 5% FBS. Both ends of the bone were cut to expose the internal bone marrow, which was flushed from both ends with a 26-G needle attached to a 10 ml syringe containing RPMI-1640 with 5% FBS into a tissue-culture treated 100-mm dish. The bone marrow cells were disaggregated by mixing with a syringe fitted with a 16-G needle. To remove remaining bone fragments, the cell suspension was passed through a 100-μm filter. The cells were centrifuged at 500 x g for 10 minutes at 4°C and counted. The cells were centrifuged again at the same settings and resuspended in sterile HBSS for retro-orbital venous injection. The cell concentration was adjusted such that each mouse received at least 5 million bone marrow cells. Retro-orbital injections were completed at least 6 h post irradiation to allow for the irradiated dying cells to clear before bone marrow reconstitution.

### Tissue processing of murine samples

Peritoneal or alveolar cells were retrieved by flushing the peritoneal cavity or the bronchoalveolar with 5 ml PBS containing 2 mM EDTA. For acute peritonitis, mice were injected intraperitoneally with 10 μg or 1 mg zymosan (Sigma cat#Z4250) 3 h before analysis. Cells were then centrifuged and resuspended in flow buffer (PBS containing 1% BSA and 2 mM EDTA). Blood was collected in eppendorf tubes containing 15 μl of 500 mM EDTA and red blood cells were lysed using BD PharmLyse lysis buffer. Omentum and mesentery samples were collected, washed in PBS, and subsequently digested for 45 min at 37°C using Collagenase IV (1 mg/mL) and DNAse I (100 μg/ml) diluted in PBS containing 1% BSA. Cell suspensions were further homogenized using a 1mL syringe with 21-G needle and filtered on a 70-μm cell strainer. To isolate mesothelium-associated tissue surface macrophages, peritoneal organs (visceral adipose tissue, spleen, omentum, liver and kidneys) were collected after performing peritoneal lavage, washed again in 1X PBS, pooled together in a 15-ml conical tube containing 5 ml of 0.25% Trypsin-EDTA (cat# 25200056) solution and incubated at 37°C with agitation for 10 minutes, as previously described (*49*). The resulting cell suspension was then filtered on a 70-μm cell strainer, briefly vortexed to avoid cellular adhesion and washed twice with PBS. For quantification of peritoneal cells, cells were stained with acridine orange and counted on a Nexcelom Cellometer Auto X4 cell counter. Quantification of omental and mesothelium-associated cells was based on cytometer counts provided by the Cytek Aurora spectral cytometer.

Livers were collected based on our previously published protocol. The liver was perfused with PBS through the portal vein until it turned pale and then collected. The gallbladders were discarded. Approximately 0.5g of the left lobe was weighted, minced, and transferred to collagenase- and DNase I-containing (0.75 mg/mL and 50 µg/mL, respectively) media on ice until all the organs from all the animals were harvested. Livers in enzymatic solution were placed on a rotating shaker for 30 min at 37°C for digestion. Liver digest was then passed through 70-µm cell strainer and washed with cold complete DMEM containing fetal bovine serum (FBS) to deactivate enzymatic activities. Hepatocytes were pelleted by centrifuging at 50xg, 3 min, at 4°C and discarded. The supernatant was transferred to a new 50 ml falcon tube, and non-parenchymal cells (NPC) were pelleted at 900 rpm, 7 min, at 4°C. After discarding the supernatant from the NPC pellet, red blood cells were lysed by resuspending cell pellets with 1mL of ACK lysis buffer (Corning) and incubated for 5 min at room temperature. Ten ml of PBS was then added, and cells were re-pelleted at 900 rpm, 7 min, at 4°C. This cell pellet was then used for downstream analyses such as flow cytometry.

Hearts were perfused with 10mL of cold PBS, weighed, minced, and digested for 40 min at 37 °C in 3mL of Dulbecco’s modified Eagle medium (DMEM) (Gibco, 11965-084) containing 167 µl of 4,500 U mL^−1^ collagenase IV (Millipore Sigma, C5138), 75 µl of 2,400 U mL^−1^ hyaluronidase I (Millipore Sigma, H3506), and 30 µl of 6,000 U mL^−1^ DNAse I (Millipore Sigma, D4527). Enzymes were deactivated with Hanks’ balanced salt solution containing 2% FBS and 0.2% BSA and filtered through 70 µm strainers. Cells were incubated with ammonium-chloride-potassium (ACK) lysis buffer (Gibco, A10492-01) for 3 min at room temperature. Cells were washed with DMEM and resuspended in 100 µl of FACS buffer (PBS containing 2% FBS and 2 mM EDTA). Cells were incubated in a 1:200 dilution of fluorescence-conjugated monoclonal antibodies for 30 min at 4 °C. Samples were washed with 1mL of FACS buffer and resuspended in a final volume of 300 µl FACS buffer.

### Flow cytometry analysis

For flow cytometry analysis of the peritoneal fluid, bronchoalveolar Lavage (BAL), and liver samples, cells were washed in PBS, stained with ZombieNIR (cat# 423106), washed again, and resuspended with the antibody mix diluted in flow buffer containing BD Horizon^TM^ Brilliant Stain Buffer (cat#563794) for 30 min at 4°C. Antibodies were conjugated to fluorochromes indicated in the figures. Clones used were the following : ICAM2 (3C4), CD45 (30-F11), F4/80 (BM8), CD11b (M1/70), MHC-II (M5/114.15.2), CD226 (TX42.1), LYVE1 (ALY7), XCR1 (ZET), Sirpα (P84), CD115 (AFS98), Ly6G (1A8), Ly6C (HK1.4), CD11c (N418), CD206 (C068C2), CD73 (TY/11.8), CD62P (RB40.34), MerTK (108928), CD45.1 (A20), CD45.2 (104), FRB (10/FR2), CD169 (3D6.112), CD64 (X54-5/7.1), CD19 (1D3).

For analysis of dendritic cells, monocytes, neutrophils, and macrophages in the heart, the antibodies used were BV421 anti-mouse CCR2 (Biolegend, 150605), BV510 anti-mouse Ly-6C (BioLegend, 128033), BV605 anti-mouse CD11b (BioLegend, 101257), BV711 anti-mouse I-A/I-E (BioLegend, 107643), BV785 anti-mouse CD45.2 (BioLegend, 109839), Spark Blue 550 anti-mouse Ly-6G (BioLegend, 127664), PerCP-Cyanine5.5 anti-mouse CD45 (BioLegend, 103132), PE-Dazzle 594 anti-mouse CD11c (BioLegend, 117348), PE-Cyanine7 anti-mouse/rat XCR1 (BioLegend, 148238), APC anti-mouse CD64 (BioLegend, 139306), Alexa Fluor 700 anti-mouse CD172a (BioLegend, 144022), and APC-Cyanine7 anti-mouse CD45.1 (BioLegend, 110716). All flow cytometry data were acquired on a Cytek Aurora spectral cytometer (5 lasers configuration). Analysis of flow cytometry data, including unsupervised analysis, was conducted using FlowJo.

### Statistical analysis and reproducibility

All measurements were taken from distinct samples, and the number of subjects in each experiment or analysis was clearly indicated either in the text or in figure legends. Significance was evaluated using the statistical tests indicated in each figure legend. These analyses were performed using the Prism-GraphPad software. p < 0.05 was considered statistically significant.

---

**Supplementary Figure 1.**
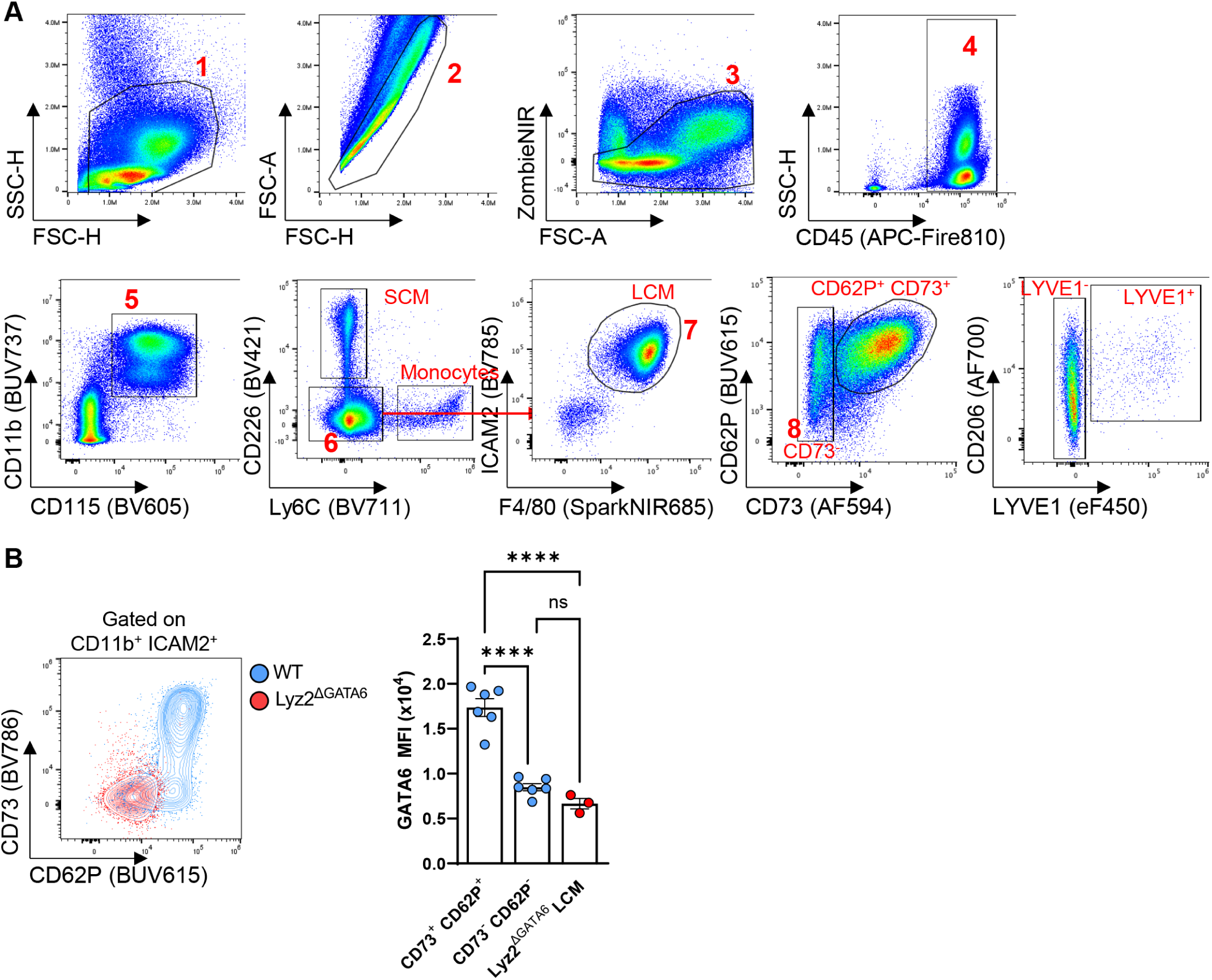
Gating strategy for peritoneal macrophage subsets and Gata6-dependent regulation of LCMs. (**A**) Gating strategy used to identify peritoneal monocytes, SCMs, and LCM subsets. (**B**) Representative flow cytometry plot of CD62P and CD73 expression (left) and quantification of Gata6 expression (right) in LCMs from WT and Lyz2^ΔGata6^ mice. Data representative of 3 independent experiments. An ANOVA was used for statistical analysis. This figure accompanies main Figure 1. ****P < 0.0001.

**Supplementary Figure 2.**
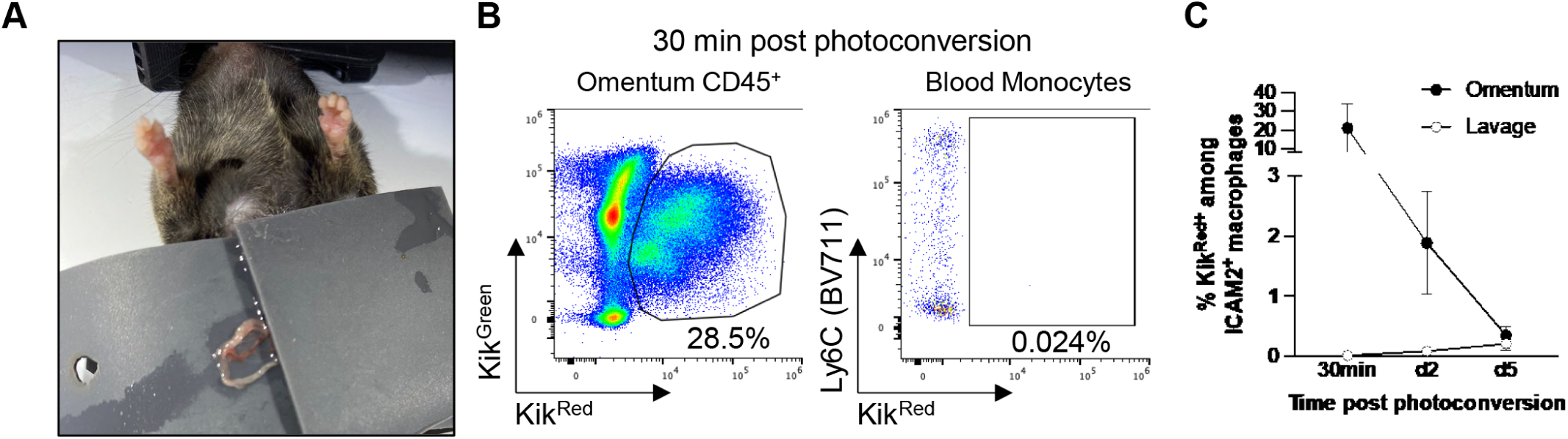
Photoconversion-based tracking of omental macrophage dynamics. (**A**) Illustration of the experimental setup enabling omentum externalization and photoconversion while shielding the peritoneal cavity. (**B**) Flow cytometry plots evaluating Kikume^Green^ and Kikume^Red^ among omental CD45^+^ cells and blood monocytes (CD11b^+^ CD115^+^) 30 minutes after photoconversion (**C**) Proportions of KikRED^+^ cells among ICAM2^+^ macrophages in the omentum and peritoneal wash following omentum photoconversion. Data were pooled from three independent experiments. This figure accompanies main Figure 2.

**Supplementary Figure 3.**
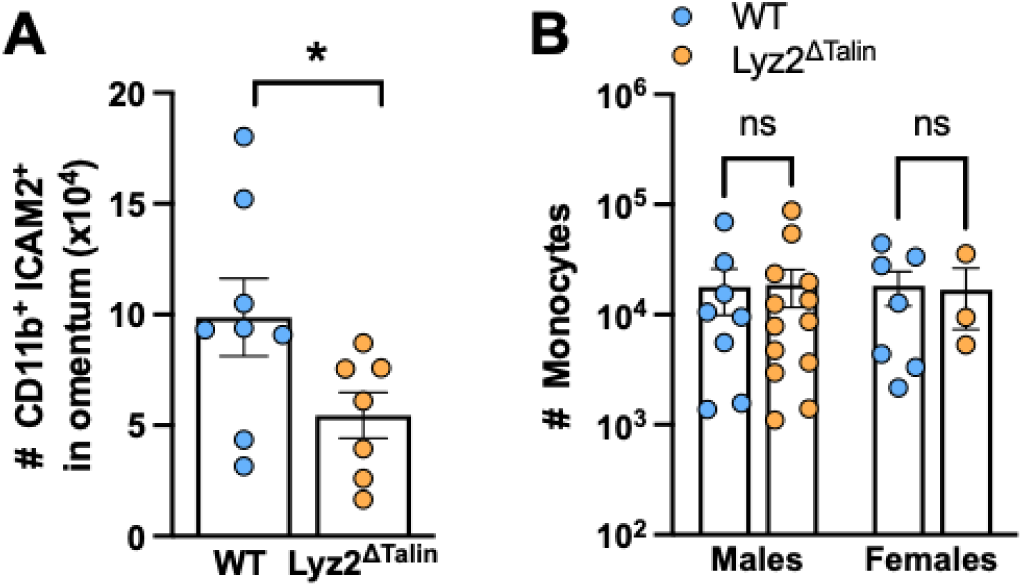
Counts of ICAM2⁺ LCMs and monocytes in talin-deficient mice versus controls across sexes following zymosan challenge. (**A**) Quantification of CD11b^+^ICAM2^+^ LCMs in omentum from WT (n=8) and Lyz2^ΔTalin^ (n=7) mice 3 hours after intraperitoneal injection of 1 mg zymosan. Data were pooled from two independent experiments. (**B**) Quantification of peritoneal monocytes from WT (male n=8, female n=7) and Lyz2^ΔTalin^ (male n=13, female n=3) mice. Data were pooled from three independent experiments. ANOVA (panel B) and Mann-Whitney (panel A) tests were used for statistical analysis. This figure accompanies main Figure 3.

**Supplementary Figure 4.**
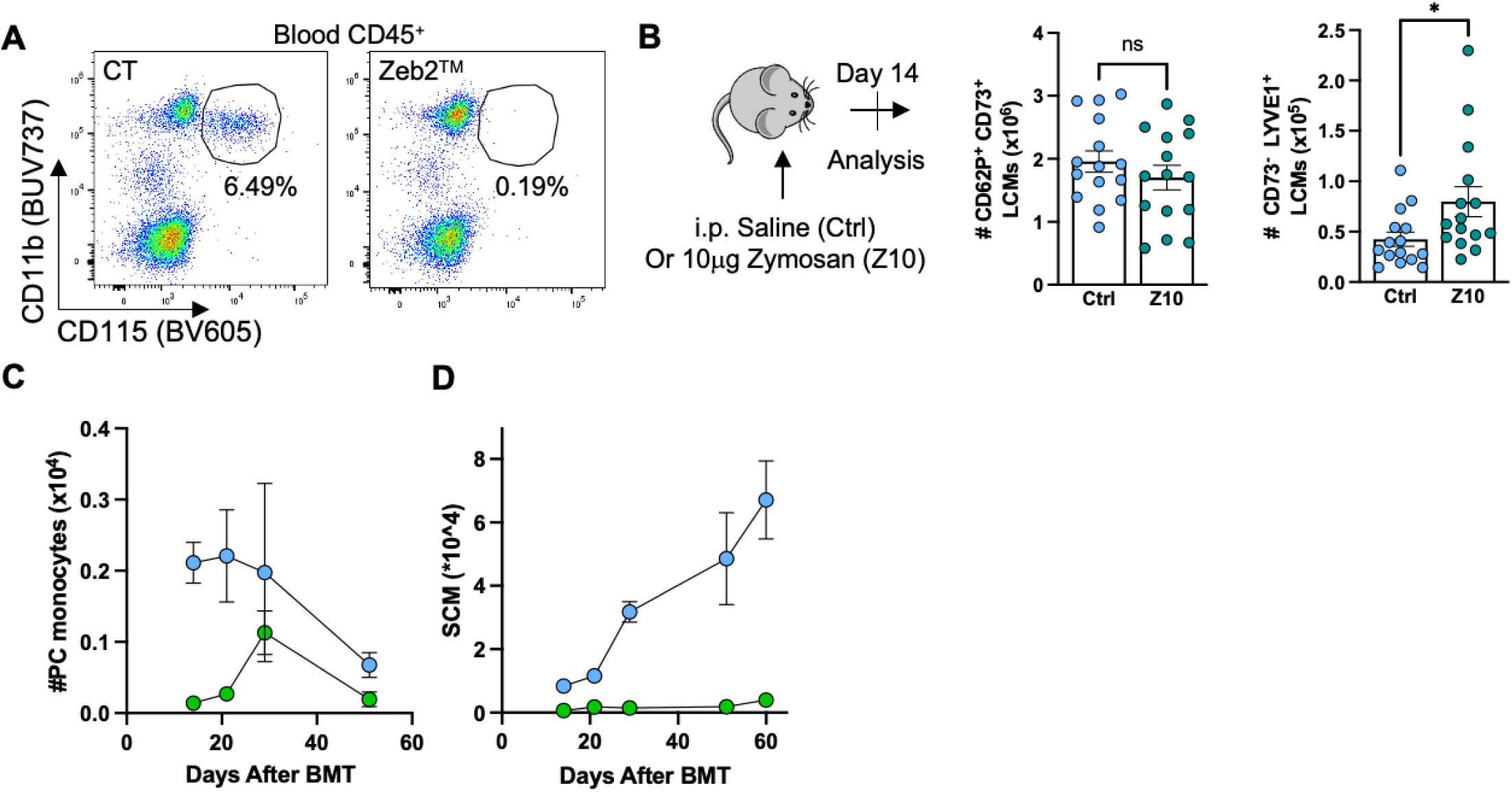
Quantification of blood and peritoneal monocytes, along with monocyte-derived peritoneal macrophage subsets, following zymosan challenge or BMT. (**A**) Flow cytometry plots representing the blood monocyte population in WT and Zeb2^TM^ mice. (**B**) Experimental scheme used to evaluate monocyte contribution to peritoneal macrophage populations following zymosan challenge (left) and quantification of peritoneal CD62P^+^ CD73^+^ and CD73^-^ LYVE1^+^ LCMs 14 days post intraperitoneal injection of saline (Ctrl, n=15) or 10 μg zymosan (Z10, n=15). Data were pooled from three independent experiments. (**C, D**) Quantification of monocytes (C) and SCMs (D) at 14 days (WT n=2, Zeb2^TM^ n=4), 21 days (WT n=3, Zeb2^TM^ n=4), 29 days (WT n=3, Zeb2^TM^ n=4), 51 days (WT n=4, Zeb2^TM^ n=3), and 60 days (WT n=9, Zeb2^TM^ n=6) post BMT. WT group (blue dots) are CD45.1 recipient mice transplanted with 50:50 Lyz2^Cre^ x R26^TdTomato^ and CCR2^gfp/gfp^ bone marrow after lethal irradiation. Zeb2^TM^ group (green dots) are CD45.1 recipient mice transplanted with Zeb2^TM^ bone marrow after lethal irradiation. Mann-Whitney tests were used for statistical analysis. This figure accompanies main Figure 5. *P<0.05.

